# Long-term dietary intervention reveals resilience of the gut microbiota despite changes in diet and weight

**DOI:** 10.1101/729327

**Authors:** Gabriela K Fragiadakis, Hannah C. Wastyk, Jennifer L. Robinson, Erica D. Sonnenburg, Justin L. Sonnenburg, Christopher D. Gardner

**Affiliations:** Microbiology & Immunology, Stanford School of Medicine, Stanford, CA; Department of Bioengineering, Stanford School of Medicine, Stanford, CA; Stanford Prevention Research Center, Department of Medicine, Stanford School of Medicine, Stanford, CA

## Abstract

With the rising rates of obesity and associated metabolic disorders, there is a growing need for effective long-term weight loss strategies, coupled with an understanding of how they interface with host physiology. While diet is a critical and promising area of focus, it has been difficult to identify diets that are broadly effective in long-term weight management. To explore the interaction between specific diets and bacteria within the gut, we tracked microbiota composition over a 12-month period as part of a larger dietary intervention study of participants consuming either a low-carbohydrate or low-fat diet. While baseline microbiota composition was not predictive of weight loss, each diet resulted in substantial changes in the microbiota three months after the start of the intervention; some of these changes were diet-specific and others tracked with weight loss. After these initial shifts, the microbiota returned near its original baseline state for the remainder of the intervention, despite participants maintaining their diet and weight loss for the entire study. These results suggest a resilience to perturbation of the microbiome’s starting state. When considering the established contribution of obese-associated microbiotas to weight gain in animal models, microbiota resilience may need to be overcome for long-term alterations to host physiology.

## Introduction

Current rates of obesity are alarmingly high and continue to increase each year,^1^ a trend that was originally confined to more affluent societies but has now begun to spread to the developing world.^2^ Diseases associated with obesity include heart disease, diabetes, and respiratory conditions, all of which contribute to lower life expectancy and quality of life. Countries faced with these trends have not been able to reverse them, despite large-scale public health and medical efforts for weight management. In order to combat these rising health concerns, and to circumvent the need for medication, many turn to diet as a way to target weight loss. However, in the context of obesity, weight modulation through diet has been variably effective and is often largely ineffective for long-term weight management. While weight reduction diets can be effective in the short term, there is evidence indicating a “memory” of obese status that contributes to post-dieting weight gain.^3–5^ The recalcitrant nature of diet effectiveness leaves many individuals at a loss for solutions, and to bear not only the burden of their health concerns, but also a misplaced sense of failure in personal responsibility that often is perpetuated by the medical community. The driving component behind ineffective long-term weight management is largely unknown, but recent studies have shown individual gut microbiota (or microbiome) signatures to be predictive of the extent of post-dieting weight gain.^6^

Previous work has established a relationship between obesity and the microbiome, including the causal role of obese-associated microbiotas to confer weight gain when transplanted into lean mice.^7–10^ When placed on the same calorically dense diet, germ-free mice have 40% lower body fat content than conventionally raised mice. Furthermore, when the distal gut microbiome of the obese mice is transferred to germ-free mice, the colonized mice experience a 60% increase in body fat within two weeks, despite no change in diet.^11^ In addition, certain microbial taxa have been shown to be associated with obesity or leanness and change in abundance during weight gain or loss.^12^ These observations may be explained by aspects of diet and the microbiota’s influence on nutritional energy extraction, which affects host fat storage in adipose tissue.^13^ Further, the microbiota has also been shown to affect intestinal permeability in obese mice thereby promoting the translocation of bacterial products and resulting in higher levels of the low-grade inflammation, a characteristic of individuals with obesity or insulin resistance.^14^ However, there is a paucity of data examining the weight loss diets and microbiome in humans.

Due to both the malleability and high degree of inter-individual variance of the microbiota, diets based on an individual’s microbiome may be a path forward in identifying effective weight loss strategies in humans. The advantage of personalized diets over universal dietary recommendations in controlling postprandial glycemic responses was recently demonstrated—an approach that used individualized aspects of the gut microbiome as parameters.^15^ While this is a promising demonstration of the predictive power of an individual’s microbiome in health management, more work is needed to better understand the interactions between specific aspects of diet and the microbiome, and the resulting effect on weight loss and weight loss maintenance.

The Diet Intervention Examining The Factors Interacting with Treatment Success (DIETFITS) clinical trial compared a healthy low-carbohydrate vs. healthy low-fat for weight loss, in a yearlong dietary intervention study.^16^ The objective was to observe how host factors, such as a metabolism-related genotype and insulin resistance, affected the success of the two diets as measured by weight loss. It was shown that while participants did lose a significant amount of weight over a period of 12 months, neither diet was universally superior and specific aspects of host genotype or insulin resistance were unable to predict diet-specific weight loss.

Here we explore another individual-specific factor, the microbiome, in diet-specific weight loss from a subset of participants in the DIETFITS trial. In this exploratory analysis of the microbiome in the DIETFITS weight loss diet study, our primary objective was to determine if baseline microbiota composition or diversity was associated with weight loss success. Our secondary objective was to examine the relationship more broadly between individual components of microbiome composition, diet, and weight. This characterization will inform the ongoing challenge of long-term weight management. Weight loss trial designs that incorporate microbiota monitoring and implement microbiota-targeting diets are a logical step toward addressing the individual and global health burden of the obesity epidemic.

## Subjects and Methods

### Study design

The detailed methods for this study were previously described.^16; 17^ Briefly, 609 generally healthy, non-diabetic participants were randomized in equal proportions in a parallel design weight loss diet study to one of two diets: healthy low-carbohydrate or healthy low-fat. Enrollment for the first participant in the first cohort started in January 2013 and follow-up for the last participant in the last cohort was completed in May 2016. Randomization was performed using an allocation sequence determined by computerized random-number generation in block sizes of 8 (4 per diet group). Participants were instructed to decrease their intake of foods that contributed the most carbohydrate or fat to their diets, respectively, until they reached the lowest level of the restricted macronutrient they could sustain over the 12 month study period. During this time, weight and food intake were tracked for each participant. Dietary intake at each time point was assessed using 3 unannounced 24-hour multiple-pass recall interviews.^16^ Data collection intervals were at pre-randomization baseline, three months, six months, and 12 months. Due to the large size of the study sample, participants were enrolled in five cohorts of approximately 120 participants per cohort, with a new cohort starting approximately once every 6 months. The analysis described here is taken from a subset of participants (n=49) from whom a complete set of stool samples were collected, all of whom were volunteers from what is referred to as “cohort 3”.^16^ Study data were collected and managed using REDCap electronic data capture tools hosted at Stanford University (Research IT grant support Stanford CTSA award number UL1 TR001085 from NIH/NCRR). REDCap (Research Electronic Data Capture) is a secure, web-based application designed to support data capture for research studies.^18^ The procedures followed were in accordance with the ethical standards of the institution or regional committee on human experimentation and that approval was obtained from the relevant committee on human subjects.

### Stool collection

Stool samples were collected at five time points over the course of the study for this cohort: pre-randomization baseline, three months, six months, nine months, and 12 months. Participants were provided with stool collection kits. Stool samples were kept in participants’ home freezers (−20°C) wrapped in ice packs, until they were transferred on ice to the research lab and stored at −80°C.

### 16S rRNA gene amplicon sequencing

DNA was extracted using the MoBio PowerSoil kit according to the Earth Microbiome Project’s protocol^19^ and amplified at the V4 region of the 16S rRNA subunit gene and 250 nt paired-end Illumina sequencing reads were generated. Data was demultiplexed using QIIME 1.8 and amplicon sequence variants (ASVs) were identified using the dada2 package in R. ASVs were assigned taxonomy using the GreenGenes database (version 13.8). Subsequent analyses including diversity analysis was performed using the phyloseq package in R. Data was rarefied to 5882 reads per sample.

### Statistical analysis

Statistical comparisons for bacterial abundances were performed using the Significance Analysis of Microarrays algorithm (SAM)^20^ using the siggenes package in R. Differences were assessed using paired participant data where appropriate. Due to missing data, we randomly selected the same number of participants from each diet based on the diet group with the lower number of samples when comparing a given time point to ensure statistical comparability. Relationships between bacterial abundances and weight were assessed using a linear mixed effect model using the nlme package in R, where participant was included as a model term to address autocorrelation of multiple samples from the same participant. P-values were adjusted for multiple hypothesis testing using a Benjamini-Hochberg correction.

## Results

### Participants decreased intake of carbohydrate or fat over time

Participants in the study were monitored for weight and dietary intake and additionally provided stool samples to enable microbiota compositional assessment over the course of the intervention. Participants (n = 49, baseline BMI 28 – 40 kg/m^2^) were randomly assigned to one of two diets, healthy low-carbohydrate or healthy low-fat, for one year (**Figure 1A**, **Supplemental Table 1**). Weight and diet data for each participant were collected pre-randomization, and at three, six, and 12 months. Stool samples for microbiome analysis were collected at the same time points.

**Figure 1.**
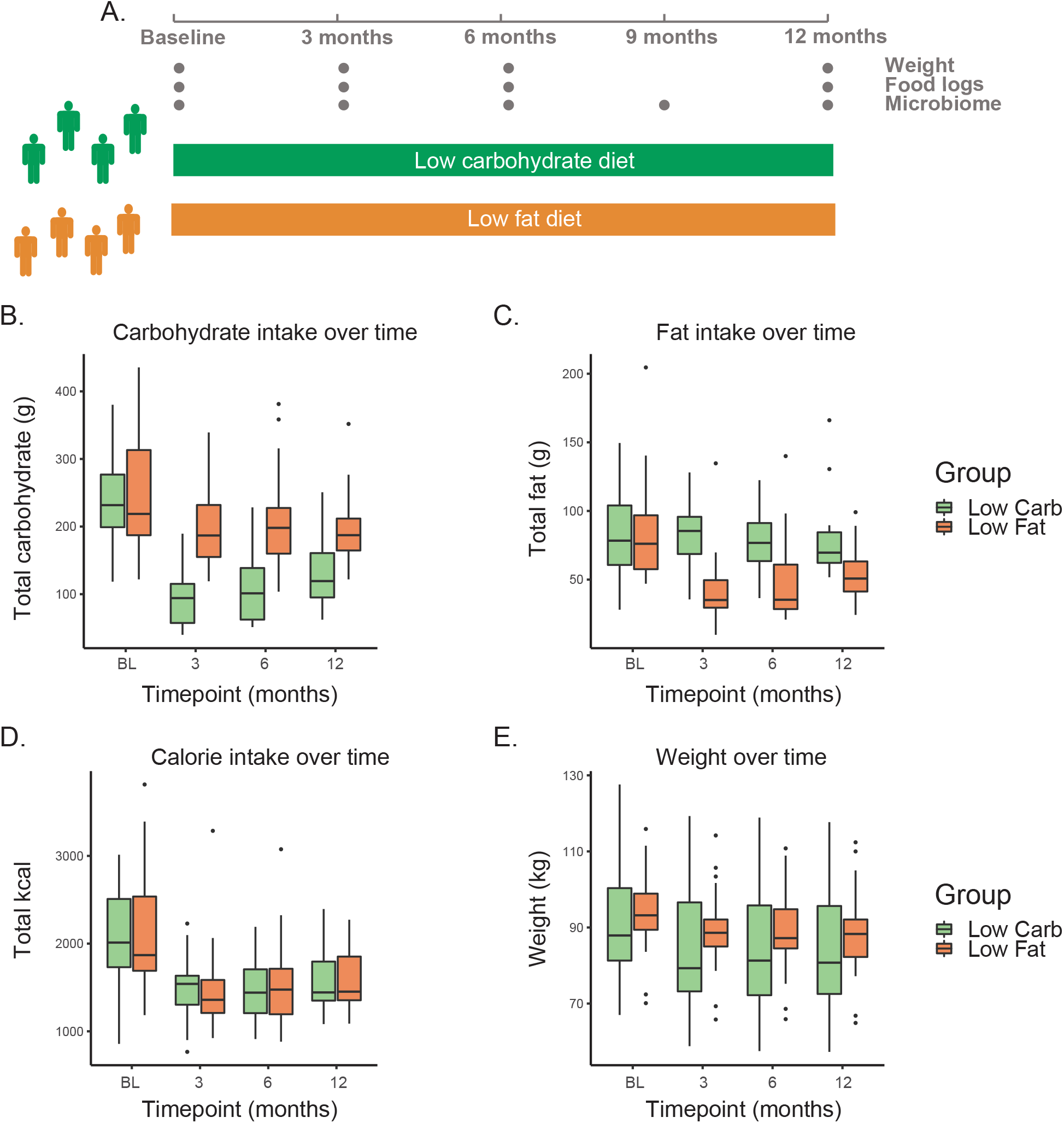
One year of low carbohydrate or low fat diet results in specific dietary alterations and weight loss. **A.** Twelve-month study design for two diet groups: healthy low carbohydrate and healthy low fat. Weight, food intake assessment, and stool for microbiome analysis were collected at the indicated sampling times. B-E. Levels of carbohydrate intake (B, grams), fat intake (C, grams), calorie intake (D, kilocalories), and weight (E, kilograms), over time for participants, separated by diet group. Green: low-carbohydrate; orange: low-fat. Significant differences indicated in **Supplemental Table 2**.

Macronutrient intake was tracked for each participant on each diet. Participants on the low-carbohydrate diet significantly decreased their carbohydrate consumption compared to baseline at every time point (**Figure 1B**, **Supplemental Table 2**). The low-fat diet participants also modestly decreased carbohydrate consumption at all time points relative to baseline. Similarly, participants in the low-fat diet significantly decreased their fat intake compared to baseline at every time point (**Figure 1C**, **Supplemental Table 2**). There was no significant change in fat intake in the low-carbohydrate diet group. Furthermore, there was no significant change in protein intake for either diet group at any time point (**Supplemental Figure 1, Supplemental Table 2**). Participants in both diets reported lowering their calorie intake by approximately 30%, and maintaining those calorie levels for the duration of the study (**Figure 1D**, **Supplemental Table 2**).

On average, both groups successfully decreased in weight over the course of 12 months (**Figure 1E**, **Supplemental Table 2**). There was no significant difference in weight loss between the two groups, with large inter-participant variance (average loss - −5.1 kg +/−6.7 for the low-carbohydrate diet and 5.6 kg +/− 5.7 sd for the low-fat diet over 12 months, **Supplemental Figure 2**). In general, weight loss was maintained over the course of the study, with a small amount of recidivism observed in the low-carbohydrate group (adjusted p-value < 0.05 between six months and 12 months, **Supplemental Figure 2**).

### Baseline microbiome composition does not predict weight loss

In addition to the factors explored in the main study, we wanted to determine whether baseline microbiota composition was an indicator of general or diet-specific weight loss. Stool samples were profiled using 16S rRNA gene amplicon sequencing, and amplicon sequence variants (ASVs) were identified using the DADA2 method ^21^ and were assigned taxonomy (Methods). We could not build a significant cross-validated model to predict weight loss from baseline microbiome composition, as described by either taxonomic summaries or by ASVs (elastic net model). Similarly, diet-specific weight loss could not be predicted from composition at baseline. In addition, no significant model could be built from composition at three months, indicating that early changes to the microbiota were not predictive of total weight loss. When participants were divided into two groups based on upper and lower quartiles of weight loss in each cohort, no significant changes in baseline composition or alpha diversity were observed. Taken together, neither baseline nor early microbiota composition was an indicator of total weight loss.

### Microbial composition exhibits resilience after initial shift

We were interested in studying the effects of dietary intervention and weight loss on the microbiome. To determine how microbial composition changed over the length of the dietary intervention, we calculated the abundance of the observed taxa at the phylum, class, order, family, and genus level for each participant at each time point. Interestingly, for each diet, we observed changes at three months relative to baseline, however these changes were not sustained throughout the remainder of the study (six months, nine months, 12 months, SAM two-class paired, **Supplemental Table 3**; specific changes are discussed later in the manuscript).^20^ The same trend was seen when testing individual ASVs despite participants maintaining their diet and lower weight throughout the course of the study (**Supplemental Table 4**).

To determine how the microbiota as a whole shifted at different time points in the study, Bray-Curtis dissimilarity was used to show the amount of shared “species” (ASVs) between samples. The first principal component of BC-distance was plotted per participant and visually showed a distinct shift at three months compared to the remaining time points (**Figure 2**, baseline shown as open circle, three months shown in red). This shift was observed in both diets, but most clearly in the low carbohydrate group, with the three-month sample being the left-most point for nearly every participant. The return of microbiota composition toward baseline status at six and 12 months occurred despite participants continuing their assigned diet and maintaining weight loss beyond the three-month period. These data suggest a resilience of the microbiota to dietary and host physiological change.

**Figure 2.**
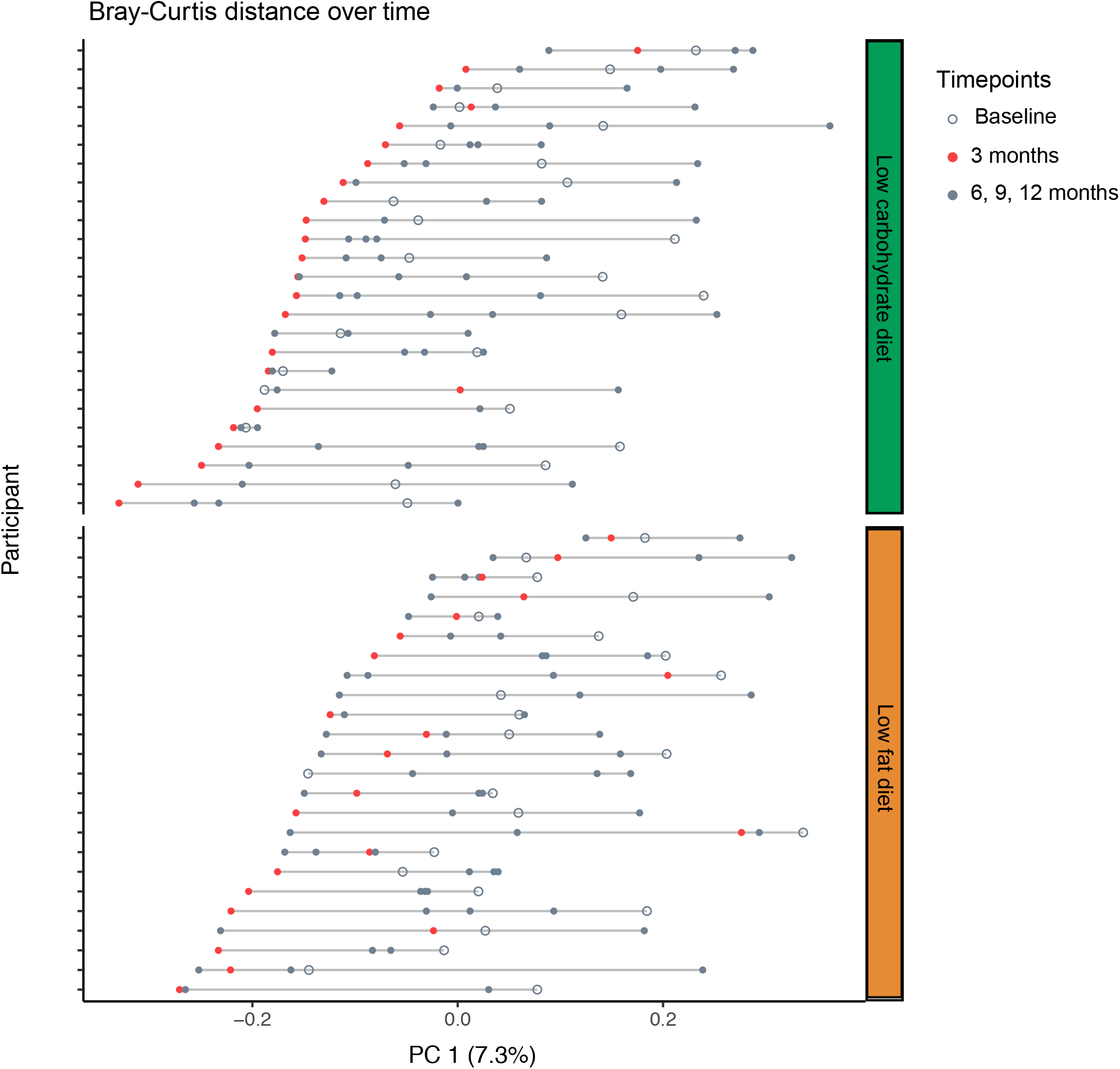
Evidence that gut microbiota composition is perturbed by diet, but exhibits resilience over time. Bray-Curtis distance between all samples was calculated, and principal coordinate analysis was used to find new axes that captured the most variance across sample distance. Values for Principal Coordinate 1 (PC1) of Bray-Curtis distance was plotted, grouped by participant and by diet. Open gray circle, baseline sample; red circle, three months sample; filled gray circle, six months, nine months, and 12 months samples.

### Each diet results in distinct changes to the microbiota during the first three months

To better understand the microbiota compositional shift that occurred at three months, we identified specific taxa that changed in relative abundance in each diet group. Each diet had a unique set of taxa that significantly changed across the cohort in the first three months. Importantly, no differences in taxa were found between the two diet groups at baseline. In the low-carbohydrate diet, changes were found in taxa from the Proteobacteria, Bacteroidetes, and Firmicutes phyla (**Figure 3A**). In the low-fat diet, changes were found in the taxa from the Actinobacteria and the Firmicutes phyla (**Figure 3B**). Several of these changes in each diet group occurred in closely related taxa, providing confidence that these were not spurious observations. Interestingly, all changes specific to the low-carbohydrate diet were increases in relative abundance, whereas all low-fat-specific changes were decreases in relative abundance; compensatory shifts in relative abundance were not uniform across the cohort. There were no observed changes in alpha (i.e. within individual) diversity in either diet, at any time point (**Supplemental Figure 3**).

**Figure 3.**
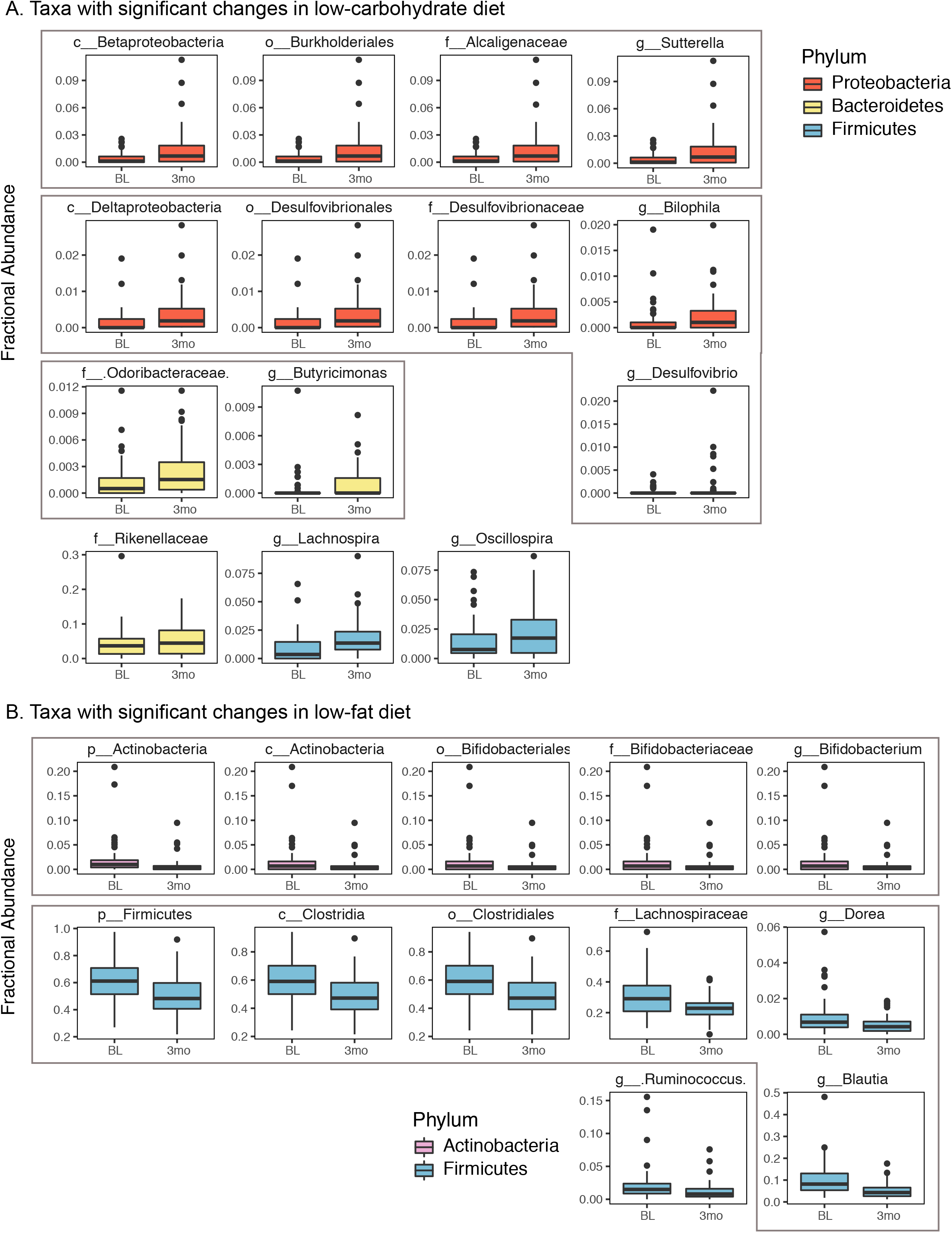
Each diet results in distinct changes in the gut microbiota composition after three-months. Diet-specific compositional changes at three months on low-carbohydrate (A) or low-fat (B) diet. Fractional abundance of taxa that significantly changed between the baseline (BL) sample and three month (3mo) sample exclusively in either diet group are shown. Significance calculated as adjusted q-value < 0.05 (SAM two-class paired). Plots colored by phylum. Gray boxes denote shared lineage. Significant changes found in both groups are shown in Figure 4A. “x_” indicates phylogenetic level where x = p, phylum; c, class; o, order; f, family; g, genus.

### Microbial compositional changes in response to both diets correlate with weight

In addition to the diet-specific changes observed at three months, several taxa that changed in relative abundance in both diets, all of which were in the Bacteroidetes phylum, which has been linked to weight loss previously^8^ (**Figure 4A**). We hypothesized that because these changes were shared by both diets, they could be due to a physiological shift in the host (i.e., weight loss) rather than a direct consequence of a differential dietary makeup. We modeled the relationship between weight and each taxonomic abundance separately for the entire study duration (confined to taxa whose abundance > 1% in at least 5% of the samples), using linear mixed effects models to account for participant autocorrelation, and assessed significance after correcting for multiple-hypothesis testing (Methods). Five of the seven taxa identified as shared changes at three months were significantly negatively associated with weight (**Figure 4B**). In addition, the genera Lachnospira and Oscillospira were negatively associated with weight, whereas, Faecalibacterium, Clostridiales, and Clostridium were positively associated with weight.

**Figure 4.**
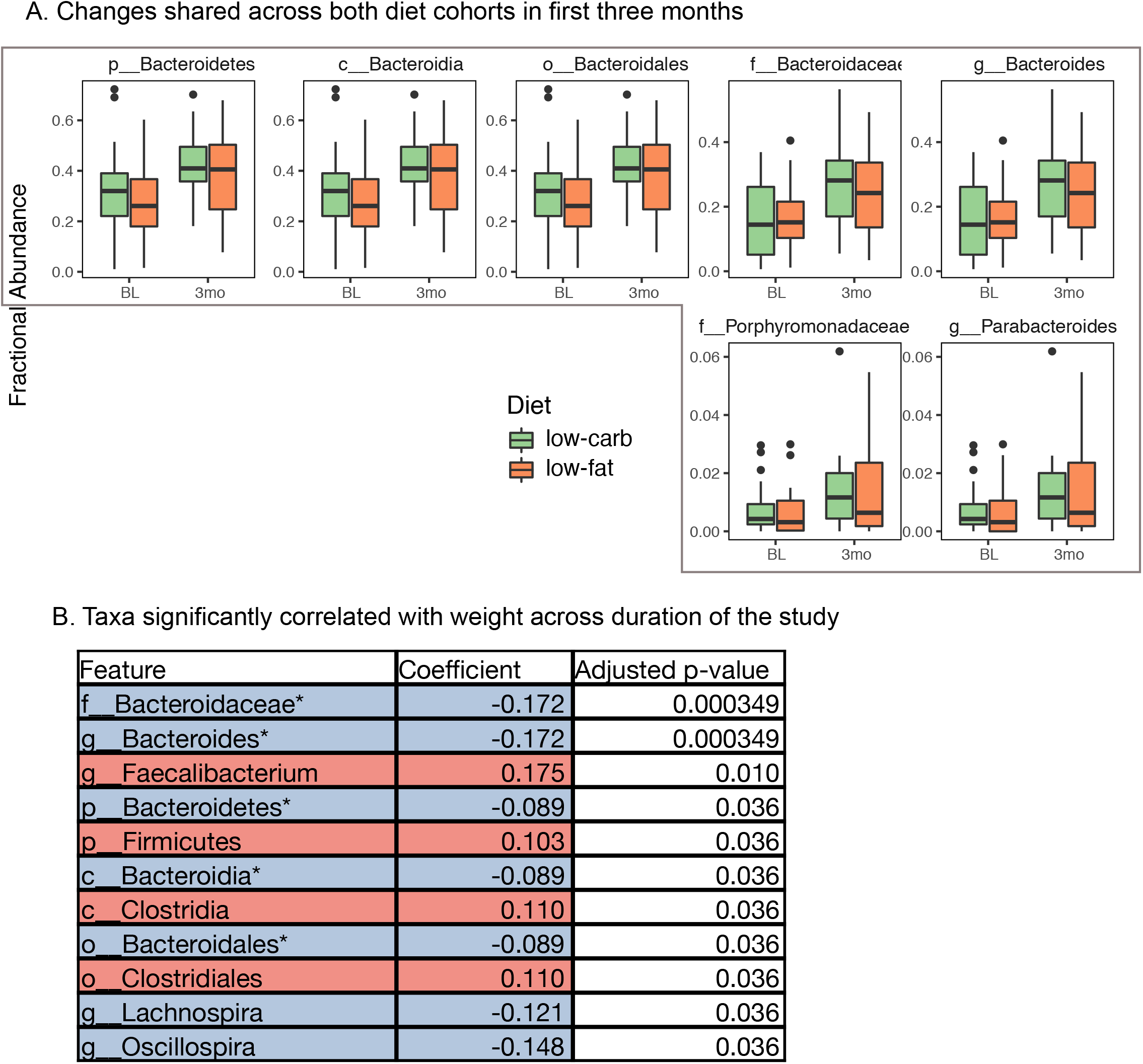
Changes observed in both diets correlate with weight. A. Compositional changes at three months shared between low-carbohydrate diet and low-fat diet. Abundance of taxa that significantly changed between the baseline sample and three month sample on both diets are shown (all participants plotted). Significance calculated as adjusted q-value < 0.05 (SAM two-class paired). Plots colored by phylum. Gray boxes denote shared phylogeny. Diet-specific changes are shown in Figure 3. “x_” indicates phylogenetic level where x = p, phylum; c, class; o, order; f, family; g, genus. B. Modeling results of taxonomic abundance and weight. Linear mixed effects models were optimized per taxa, with taxa significantly associated with weight listed in the table (adjusted p-value < 0.05, Benjamini-Hochberg). Taxa were tested whose abundance was > 1% in at least 5% of the samples. Taxa with fractional abundance positively correlated to weight are highlighted in red; taxa with fractional abundance negatively correlated to weight are highlighted in blue. * denotes taxa that were identified in Figure 4A.

## Discussion

In this work, we examine the dynamics of the microbiome in the context of weight loss and dietary change. Investigation of the microbiome is particularly crucial considering the rising rates of obesity and failure of current diet-based strategies for long-term weight loss. The original work did not find an association with either insulin resistance or genotype pattern and weight loss, leading to an interest in exploring whether other host factors predisposed certain individuals to lose more weight on specific diets. Here we used 16S rRNA gene amplicon sequencing to assess the composition of the microbiota at several intervals throughout the 12-month intervention from participants on each diet. We did not find an association between baseline microbiome composition and weight loss on either diet. This finding suggests that in our cohort, microbiome composition does not dictate the degree of weight loss on a low-carbohydrate versus low-fat a diet, nor weight loss more generally. This conclusion is in contrast to recent work that found personalized glycemic responses to specific diets, which could be predicted in part by microbiome composition^15^. This disparity could be because our studied diets, a general healthy low-carbohydrate and healthy low-fat diet, were not selected a priori as diets to specifically target the microbiome. Notably, though both examined the relationship between the microbiome and host health, each study used different outcome measures (glycemic responses versus weight loss), which may not be perfect correlates. In addition, study size and type of data are important factors in the sensitivity of these types of analyses, so it is possible that larger studies or different data types will reveal microbiome predictors of weight loss.

While we did not find an association between baseline microbiome composition and weight loss, we did observe high variance in weight loss among individuals in the study, suggesting that there are likely host factors that predispose individuals to weight loss that have yet to be elucidated. We did not find a statistically significant relationship between the degree of calorie restriction and the amount of weight loss (Supplemental Figure 4). Interestingly, this could suggest that more nuanced diets rather than calorie restriction may be important in successful, sustained weight reduction. However, it could also be that a true relationship that exists between calorie restriction and weight loss was masked by inaccuracies in self-reported diet data, which are well-established.^22^ This observation presents an opportunity for personalized diets that better suit an individual’s starting state, though substantial investigation is required to study both relevant host factors and aspects of diet that can be used as levers to impact host physiology.

While we did not find microbiome signatures predictive of weight loss, we did observe substantial changes to the microbiome during the intervention. As we delved into these responses in microbiome composition, we found that there were two forces affecting the microbiome: the change in diet (i.e., specific changes in energy and macronutrient availability), and the host physiological changes associated with weight loss. It is difficult to disentangle these effects since they co-occur. However, mouse models have shown obesity-independent effects of a high-fat diet on the microbiome, demonstrating that these forces (host physiology and diet) can exert separate and independent effects on the microbiota.^23^ We identified a set of changes in bacterial taxa during the first three months of the intervention that included both a subset that were shared between the two diets, and a subset that were only observed in one diet. On the low-carbohydrate diet, participants had increases in the relative abundance of taxa in the Proteobacteria, Bacteroidetes, and Firmicutes phyla. In contrast, participants on the low-fat diet exhibited decreases in Actinobacteria and Firmicutes. This result is consistent with previous work that observed an obesity-independent increase in Firmicutes in mice fed a high-fat diet.^23^

Among the changes that were shared between the two diets was an increase in Bacteroidetes relative abundance. Lower Bacteroidetes abundance has been observed in obese relative to lean mice and humans^7; 8^. In addition, a small cohort (n = 12) of individuals on either a carbohydrate- or fat-restricted diet exhibited an increase in Bacteroidetes abundance that correlated with weight loss^8^. Our hypothesis that these shared changes in both diets is due to weight loss rather than specific dietary changes was supported by our finding that these taxa correlated with weight throughout the study. Oscillospira and Lachnospira abundances similarly decreased with weight, consistent with a reported association of Oscillospira with leanness^12^ and lower levels of Lachnospira in obesity.^24^ In contrast, Firmicutes and members of its lineage were positively associated with weight, which is consistent with the Firmicutes:Bacteroidetes ratio being higher in obese individuals^8^.

The changes in the microbiota described here were confined to the first three months of the dietary intervention, and while there was a global directional shift in overall microbiota composition during that time, it was not sustained during later sampling times. This was despite participants largely maintaining their weight loss, and maintaining statistically and clinically meaningful dietary changes relative to baseline^16^. The microbiota exhibited resilience after the initial perturbation of diet change and weight loss, perhaps due to host physiological factors that exert a homeostatic corrective force on the microbial community to return to a long-established state. Interestingly, previous work identified a microbiome-based “memory” of obesity, which may explain the phenomenon of individuals regaining lost weight despite maintaining a previously effective diet^6^. The observed resilience described here may be an instance of the same phenomenon, and further work is needed to explore strategies to overcome this microbial resilience in cases where the microbiota is in an unhealthy state.

This study has several limitations. Our sampling of the microbiota was confined to three-month intervals, which restricts our understanding of the full dynamics of microbiome composition. While we see changes that occur at three months post-diet intervention that are no longer present at six months, it is not possible to determine when those changes first began nor when they start to disappear, as well as whether there were further oscillations not captured. In addition, these diets were not specifically designed to target the microbiota but rather were selected based on their prevalence as diets for weight loss. Future studies that assess the effect of diets that have been shown to modulate the microbiota, such as those high in dietary fiber,^25; 26^ may shed further light on the interaction between diet, the microbiome, and weight loss. Finally, while we assess weight as a primary outcome, we do not profile other elements of host health such as inflammation and immune status, which are likely affected by diet and the microbiome.^27^

In light of the rising rates of obesity and the accompanying morbidities, along with our current failures thus far in reversing such trends, there is a moral imperative to identify long-term effective solutions for weight loss. Though diet-based strategies are promising, the limited long-term efficacy points to gaps in our understanding of the interplay between specific diets and host factors, such as inflammatory and metabolic state, and the microbiome. This work identifies distinct but transient effects of diet and of host physiological state on the gut microbiota in humans. Future work toward understanding mechanisms of microbiota resilience, in the context of both microbial functionality and interaction with factors in the host such as inflammation, are needed. Next steps should include investigating the importance of an individual’s gut microbes when defining and implementing personalized diets for weight loss.

## Acknowledgements

G.K.F. and H.C.W. performed the data analysis and wrote the manuscript. J.L.R. generated data and edited the manuscript. E.D.S. advised analysis and edited the manuscript. C.D.G. and J.L.S. designed the study, advised analysis, edited the manuscript, and have primary responsibility for the final content. A thanks to Erin Avery and Audrey Southwick for data generation.

## Abbreviations

BL: baseline
ASV: amplicon sequence variant
SAM: significance analysis of microarrays
low-carb: low-carbohydrate

## Clinical trial registry

clinicaltrials.gov identifier: NCT01826591

