## Supplementary data for "Long-term dietary intervention reveals resilience of the gut microbiota despite changes in diet and weight"

Supplemental Table 1: Demographics of Participants.

|  | **Low-carbohydrate** | **Low-fat** |
| --- | --- | --- |
| **Sex, No (%)** |  |  |
| Male | 5 (20) | 5 (21) |
| Female | 20 (80) | 19 (79) |
| **Age, Mean (SD), y** | 42.6 (5.8) | 39.2 (5.5) |
| **Highest level of education achieved, No. (%)** |  |  |
| High school graduate | 1 (4) | 0 (0) |
| Some college | 5 (20) | 1 (4) |
| College graduate | 12 (48) | 9 (38) |
| Some post-graduate school | 1 (4) | 3 (13) |
| Post-graduate degree | 6 (24) | 11 (46) |
| **Race/ethnicity, No. (%)** |  |  |
| White | 20 (80) | 18 (75) |
| Hispanic | 3 (12) | 3 (13) |
| Asian | 2 (8) | 1 (4) |
| Other | 0 (0) | 2 (8) |
| **Weight, mean (SD), kg** |  |  |
| Male | 100 (14.7) | 99.7 (12.9) |
| Female | 87.1 (14.7) | 92.4 (9.82) |
| Both sexes | 89.7 (15.4) | 93.9 (10.7) |
| **Body mass index, mean (SD)** |  |  |
| Male | 34.3 (3.4) | 34.0 (3.9) |
| Female | 32.4 (4.0) | 33.6 (3.5) |
| Both sexes | 32.8 (3.9) | 33.7 (3.5) |

Supplemental Table 2: P-values for changes in weight, calories, carbohydrate, fat, and protein between two specified timepoints. Highlight shows adjusted p-values < 0.05 (paired Wilcoxon test). BL: baseline; 3mo: 3 months; 6mo, 6 months; 12mo, 12 months.

| Group | Measurement | Timepoint 1 | Timepoint 2 | Adjusted p-value |
| --- | --- | --- | --- | --- |
| Low Fat | Weight | BL | 3mo | 7.761E-05 |
| Low Fat | Weight | BL | 6mo | 9.100E-05 |
| Low Fat | Weight | BL | 12mo | 1.845E-03 |
| Low Fat | Weight | 3mo | 6mo | 7.125E-02 |
| Low Fat | Weight | 3mo | 12mo | 2.645E-01 |
| Low Fat | Weight | 6mo | 12mo | 5.034E-01 |
| Low Carbohydrate | Weight | BL | 3mo | 7.153E-07 |
| Low Carbohydrate | Weight | BL | 6mo | 7.761E-05 |
| Low Carbohydrate | Weight | BL | 12mo | 1.278E-03 |
| Low Carbohydrate | Weight | 3mo | 6mo | 7.369E-02 |
| Low Carbohydrate | Weight | 3mo | 12mo | 2.912E-02 |
| Low Carbohydrate | Weight | 6mo | 12mo | 1.757E-04 |
| Low Fat | Carbohydrate | BL | 3mo | 1.901E-02 |
| Low Fat | Carbohydrate | BL | 6mo | 4.054E-02 |
| Low Fat | Carbohydrate | BL | 12mo | 3.262E-02 |
| Low Fat | Carbohydrate | 3mo | 6mo | 7.493E-01 |
| Low Fat | Carbohydrate | 3mo | 12mo | 7.540E-01 |
| Low Fat | Carbohydrate | 6mo | 12mo | 7.493E-01 |
| Low Carbohydrate | Carbohydrate | BL | 3mo | 7.153E-07 |
| Low Carbohydrate | Carbohydrate | BL | 6mo | 4.530E-06 |
| Low Carbohydrate | Carbohydrate | BL | 12mo | 2.146E-06 |
| Low Carbohydrate | Carbohydrate | 3mo | 6mo | 8.464E-02 |
| Low Carbohydrate | Carbohydrate | 3mo | 12mo | 8.179E-03 |
| Low Carbohydrate | Carbohydrate | 6mo | 12mo | 2.260E-01 |
| Low Fat | Fat | BL | 3mo | 2.861E-06 |
| Low Fat | Fat | BL | 6mo | 5.007E-06 |
| Low Fat | Fat | BL | 12mo | 2.207E-03 |
| Low Fat | Fat | 3mo | 6mo | 2.373E-01 |
| Low Fat | Fat | 3mo | 12mo | 8.179E-03 |
| Low Fat | Fat | 6mo | 12mo | 2.373E-01 |
| Low Carbohydrate | Fat | BL | 3mo | 7.974E-01 |
| Low Carbohydrate | Fat | BL | 6mo | 3.361E-01 |
| Low Carbohydrate | Fat | BL | 12mo | 5.890E-01 |
| Low Carbohydrate | Fat | 3mo | 6mo | 3.273E-01 |
| Low Carbohydrate | Fat | 3mo | 12mo | 2.590E-01 |
| Low Carbohydrate | Fat | 6mo | 12mo | 9.168E-01 |

| Low Fat | Calories | BL | 3mo | 6.294E-05 |
| --- | --- | --- | --- | --- |
| Low Fat | Calories | BL | 6mo | 3.934E-05 |
| Low Fat | Calories | BL | 12mo | 1.207E-02 |
| Low Fat | Calories | 3mo | 6mo | 3.932E-01 |
| Low Fat | Calories | 3mo | 12mo | 3.664E-02 |
| Low Fat | Calories | 6mo | 12mo | 3.932E-01 |
| Low Carbohydrate | Calories | BL | 3mo | 1.817E-04 |
| Low Carbohydrate | Calories | BL | 6mo | 6.507E-04 |
| Low Carbohydrate | Calories | BL | 12mo | 4.802E-03 |
| Low Carbohydrate | Calories | 3mo | 6mo | 8.996E-01 |
| Low Carbohydrate | Calories | 3mo | 12mo | 3.932E-01 |
| Low Carbohydrate | Calories | 6mo | 12mo | 3.932E-01 |
| Low Fat | Protein | BL | 3mo | 2.106E-01 |
| Low Fat | Protein | BL | 6mo | 2.106E-01 |
| Low Fat | Protein | BL | 12mo | 3.559E-01 |
| Low Fat | Protein | 3mo | 6mo | 9.220E-01 |
| Low Fat | Protein | 3mo | 12mo | 9.744E-01 |
| Low Fat | Protein | 6mo | 12mo | 9.220E-01 |
| Low Carbohydrate | Protein | BL | 3mo | 9.220E-01 |
| Low Carbohydrate | Protein | BL | 6mo | 9.220E-01 |
| Low Carbohydrate | Protein | BL | 12mo | 9.888E-01 |
| Low Carbohydrate | Protein | 3mo | 6mo | 4.737E-01 |
| Low Carbohydrate | Protein | 3mo | 12mo | 2.106E-01 |
| Low Carbohydrate | Protein | 6mo | 12mo | 9.220E-01 |

Supplemental Table 3: Diet-specific differences in taxa identified between the two specified timepoints, paired by participant (significance analysis of microarrays, two class-paired, q-value < 0.05)

| Taxa | Esti q-value | Diet | Comparison |
| --- | --- | --- | --- |
| g__Blautia | 0.000 | Low-fat | BL v. 3 months |
| f__Lachnospiraceae | 0.000 | Low-fat | BL v. 3 months |
| p__Firmicutes | 0.021 | Low-fat | BL v. 3 months |
| c__Clostridia | 0.021 | Low-fat | BL v. 3 months |
| o__Clostridiales | 0.021 | Low-fat | BL v. 3 months |
| g__Bacteroides | 0.021 | Low-fat | BL v. 3 months |
| f__Bacteroidaceae | 0.021 | Low-fat | BL v. 3 months |
| g__Dorea | 0.021 | Low-fat | BL v. 3 months |
| g__Parabacteroides | 0.021 | Low-fat | BL v. 3 months |
| f__Porphyromonadaceae | 0.021 | Low-fat | BL v. 3 months |
| o__Bacteroidales | 0.022 | Low-fat | BL v. 3 months |
| c__Bacteroidia | 0.022 | Low-fat | BL v. 3 months |
| p__Bacteroidetes | 0.022 | Low-fat | BL v. 3 months |
| p__Actinobacteria | 0.022 | Low-fat | BL v. 3 months |
| g__.Ruminococcus. | 0.023 | Low-fat | BL v. 3 months |
| c__Actinobacteria | 0.023 | Low-fat | BL v. 3 months |
| g__Bifidobacterium | 0.023 | Low-fat | BL v. 3 months |
| f__Bifidobacteriaceae | 0.023 | Low-fat | BL v. 3 months |
| o__Bifidobacteriales | 0.023 | Low-fat | BL v. 3 months |
| g__Lachnospira | 0.000 | Low-carb | BL v. 3 months |
| g__Bacteroides | 0.000 | Low-carb | BL v. 3 months |
| f__Bacteroidaceae | 0.000 | Low-carb | BL v. 3 months |
| f__Desulfovibrionaceae | 0.000 | Low-carb | BL v. 3 months |
| o__Desulfovibrionales | 0.000 | Low-carb | BL v. 3 months |
| c__Deltaproteobacteria | 0.000 | Low-carb | BL v. 3 months |
| f__Rikenellaceae | 0.007 | Low-carb | BL v. 3 months |
| g__Butyricimonas | 0.022 | Low-carb | BL v. 3 months |
| g__Oscillospira | 0.022 | Low-carb | BL v. 3 months |
| g__Bilophila | 0.022 | Low-carb | BL v. 3 months |
| f__.Odoribacteraceae. | 0.022 | Low-carb | BL v. 3 months |
| g__Parabacteroides | 0.022 | Low-carb | BL v. 3 months |
| f__Porphyromonadaceae | 0.022 | Low-carb | BL v. 3 months |
| p__Bacteroidetes | 0.023 | Low-carb | BL v. 3 months |
| o__Bacteroidales | 0.023 | Low-carb | BL v. 3 months |
| c__Bacteroidia | 0.023 | Low-carb | BL v. 3 months |
| o__Burkholderiales | 0.030 | Low-carb | BL v. 3 months |
| c__Betaproteobacteria | 0.030 | Low-carb | BL v. 3 months |
| g__Sutterella | 0.030 | Low-carb | BL v. 3 months |
| f__Alcaligenaceae | 0.030 | Low-carb | BL v. 3 months |
| g__Desulfovibrio | 0.038 | Low-carb | BL v. 3 months |
| g__Ruminococcus | 0.034 | Low-carb | BL v. 6 months |

Supplemental Table 4: Diet-specific differences in Amplicon Sequence Variants (ASVs) identified between the two specified timepoints, paired by participant (significance analysis of microarrays, two class-paired, q-value < 0.05)

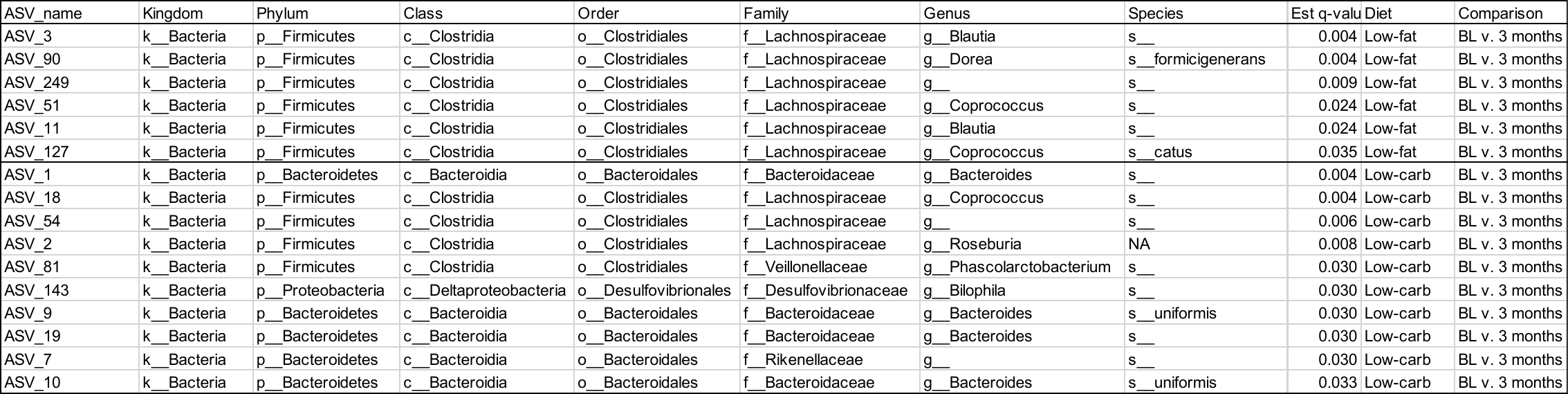

Supplemental Figure 1: Protein intake level in grams over time for participants, separated by diet group. Green: low-carbohydrate; orange: low-fat. Significant differences indicated in Supplemental Table 2.

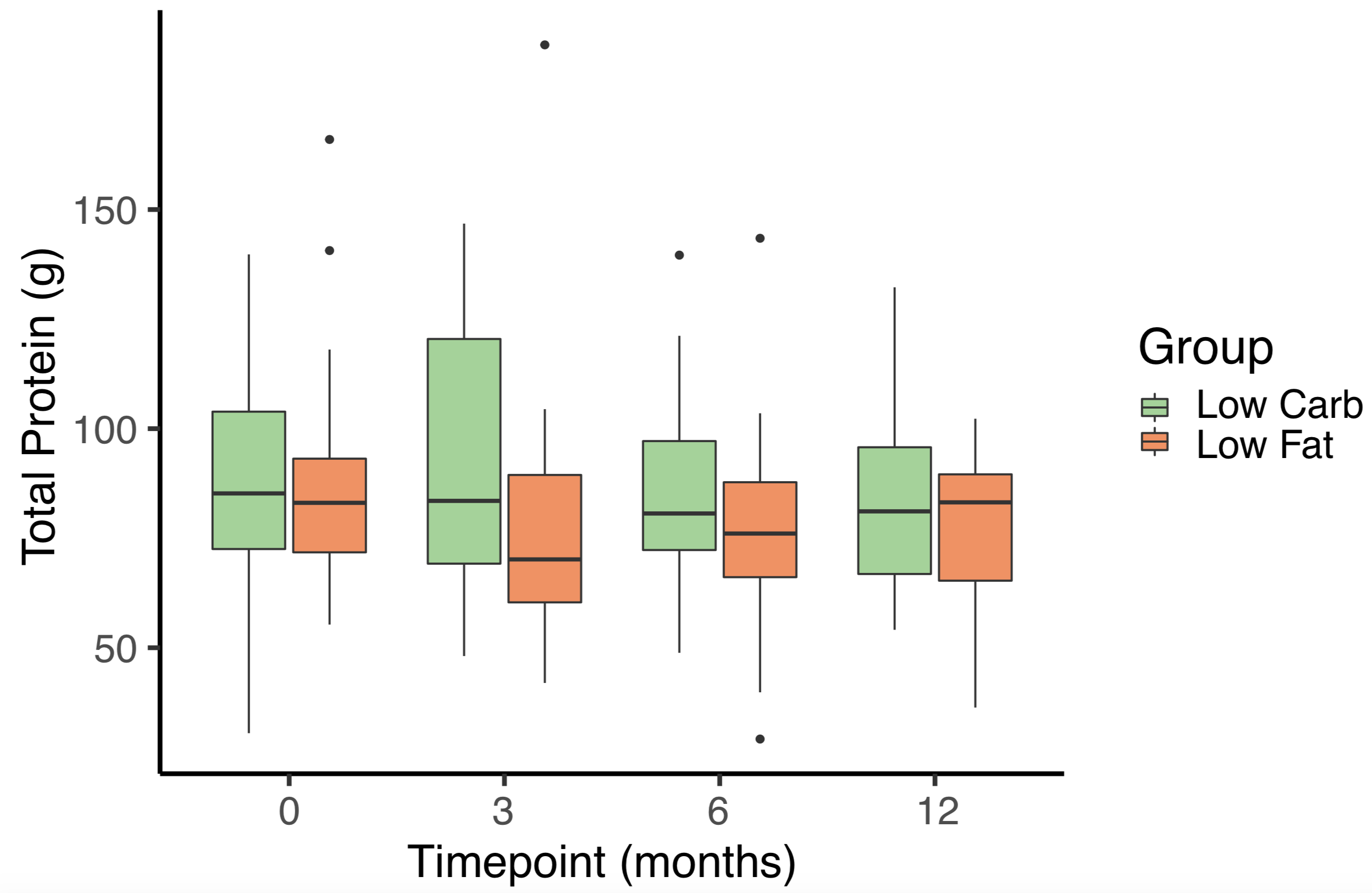

Supplemental Figure 2: Distribution of weight loss in participants by diet. Participants in each diet group were ordered based on their change in weight between baseline and 12 months. Weights shown in kilograms. Green, low-carbohydrate group; orange, low-fat group.

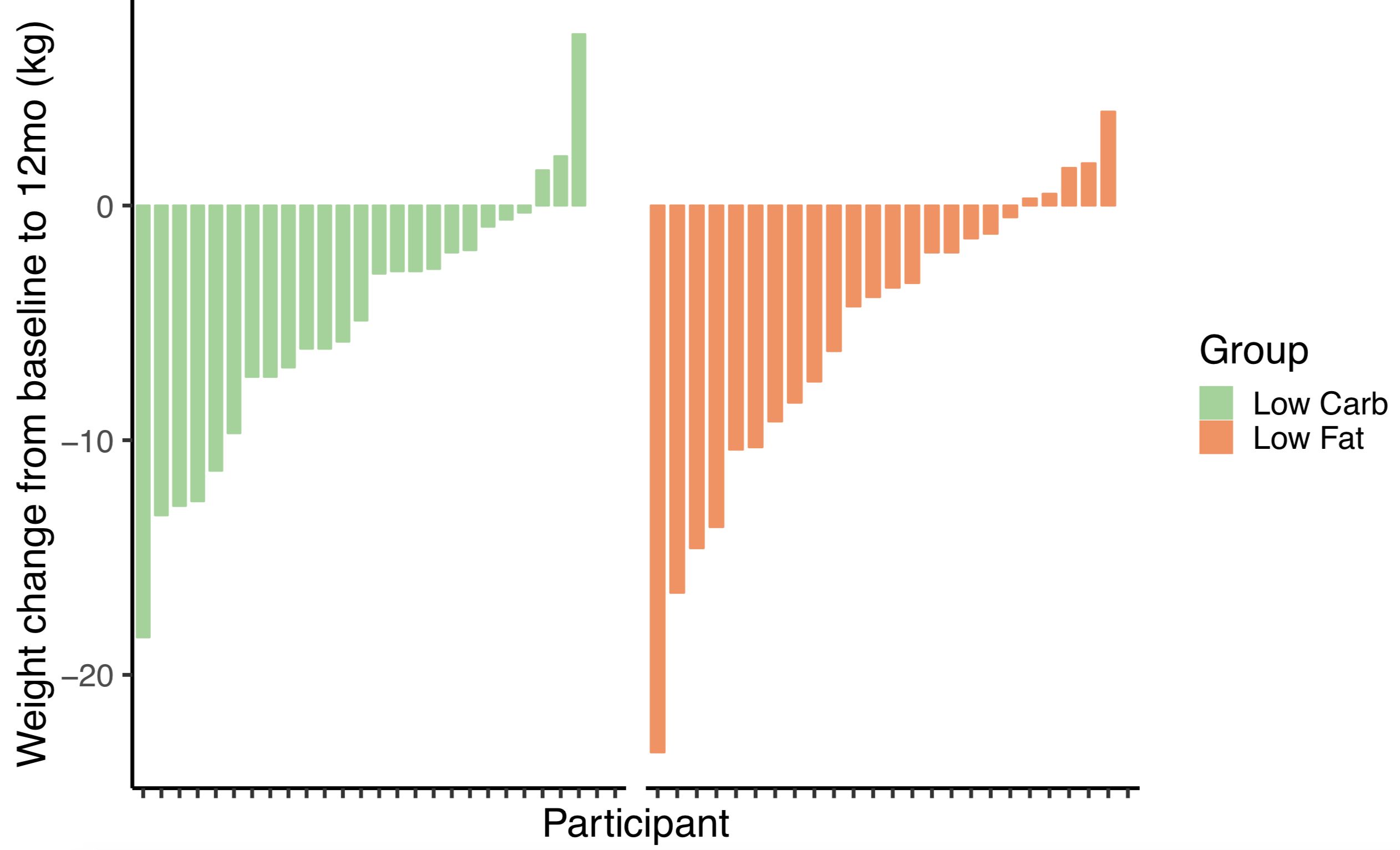

Supplemental Figure 3: Alpha diversity over time as measured by the number of observed amplicon sequence variants (ASVs), separated by diet group. Green, low-carbohydrate group; orange, low-fat group.

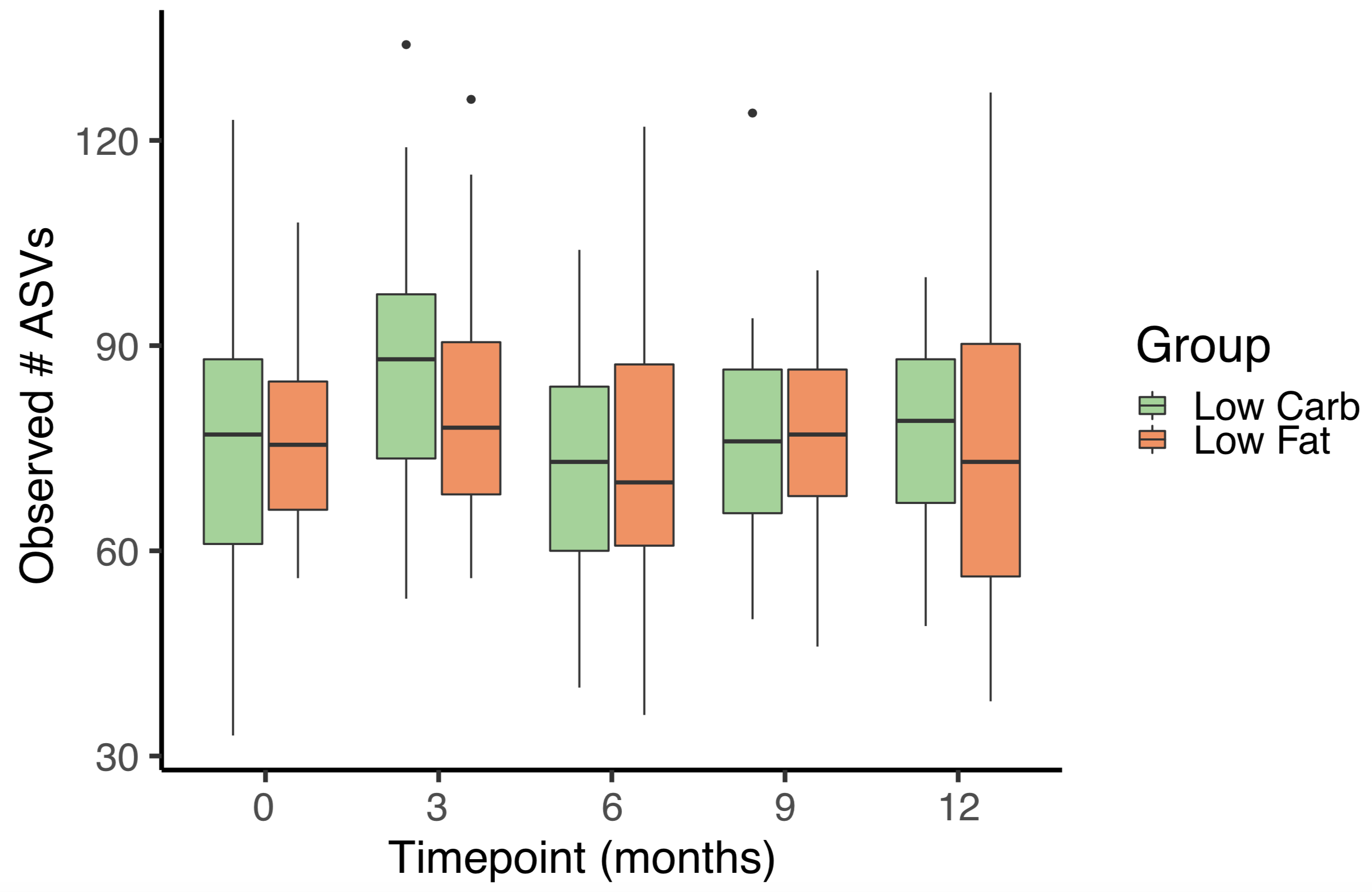

Supplemental Figure 4: Correlation between weight loss from baseline to 12 months in kilograms and calorie change from baseline to 12 months in kilocalories per day. These metrics did not significantly correlate (p > 0.05).

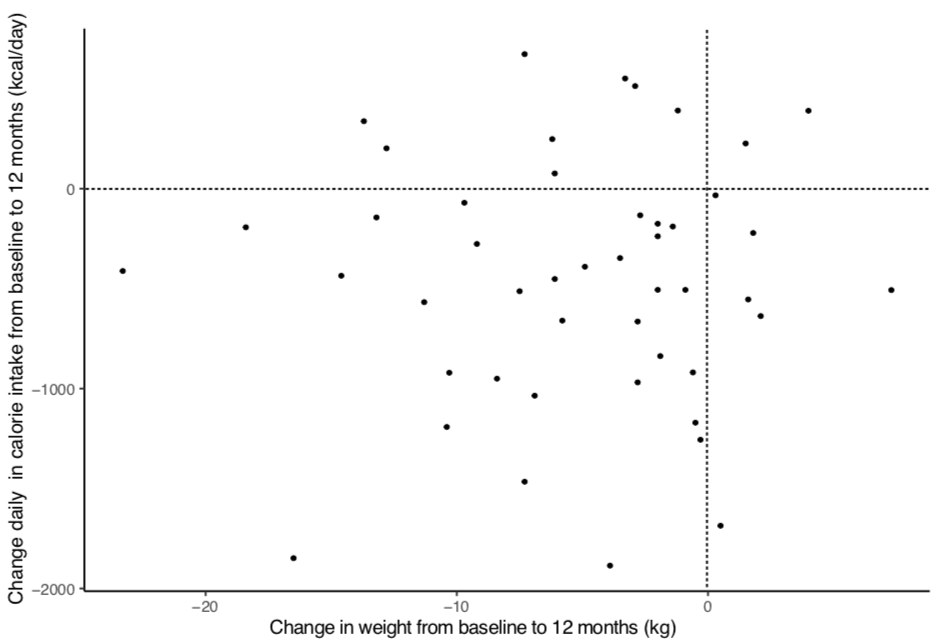
